# Unmasking the cryptic immunopeptidome of EZH2 mutated diffuse large B-cell lymphomas through combination drug treatment

**DOI:** 10.1101/2021.09.01.458572

**Authors:** Christopher M. Bourne, Sung Soo Mun, Tao Dao, Zita E. H. Aretz, Ron S. Gejman, Andrew Daman, Katsuyoshi Takata, Christian Steidl, Martin G. Klatt, David A. Scheinberg

## Abstract

Exploring the repertoire of peptides presented on major histocompatibility complexes (MHC) has been utilized to identify targets for immunotherapy in many hematological malignancies. However, such data have not been described systematically for diffuse large B-cell lymphomas (DLBCL), which might be explained by the profound downregulation of MHC expression in many DLBCLs, and in particular in the EZH2-mutated subgroup. Epigenetic drug treatment, especially in the context of interferon gamma (IFNg), restored MHC expression in DLBCL. DLBCL MHC-presented peptides were identified via mass spectrometry following tazemetostat or decitabine treatments alone, or in combination with IFNg. Such treatment synergistically increased MHC class I surface protein expression up to 50-fold and class II expression up to 3-fold. Peptides presented on MHC complexes increased to a similar extent for MHC class I and remained constant for class II. Overall, these treatments restored the diversity of the immunopeptidome to levels described in healthy B cells and allowed the systematic search for new targets for immunotherapy. Consequently, we identified multiple MHC ligands from *regulator of G protein signaling 13* (RGS13) and *E2F transcription factor 8* (E2F8) on different MHC alleles, none of which have been described in healthy tissues and therefore represent tumor-specific MHC ligands, which are unmasked only after drug treatment. Overall, our results show that EZH2 inhibition in combination with decitabine and IFNg can expand the repertoire of MHC ligands presented on DLBCLs by revealing cryptic epitopes, thus allowing the systematic analysis and identification of new potential immunotherapy targets.

**Key points:**

- Combination therapy of interferon gamma with epigenetic regulators leads to large increases in the immunopeptidome of DLBCL.
- HLA ligands from proteins RGS13 and E2F8 may provide DLBCL-specific targets for immunotherapy.

## Introduction

Diffuse Large B cell Lymphoma (DLBCL) is the most common lymphoma type in the western hemisphere. About 60% of patients can be cured of DLBCL using the standard chemo-immunotherapy, R-CHOP, but successful treatment remains challenging in relapsed patients^1^. As the R-CHOP regimen can cause considerable toxicity, which is poorly tolerated by older patients^2^, therapeutic agents that minimize adverse side effects while still demonstrating anti-tumor efficacy are attractive. Immunotherapy has seen remarkable anti-tumor efficacy and on-tumor specificity ^3–6^ and identification of neoepitopes that are specific to cancer cells can maximize on-tumor efficacy while minimizing off-target effects on healthy tissue ^7–9^. A number of cell surface targets for antibody therapies now exist^10^. Identification of suitable targets for T cell immunotherapy relies on immunoprecipitation of MHC complexes and subsequent analysis of the bound peptides via mass spectrometry (MS), which has been performed on solid tumors, leukemias, and myelomas^11–16^.

In contrast, no systematic descriptions of the immunopeptidome of DLBCL are available, possibly due to the ability of DLBCL to downregulate antigen presentation and evade immune recognition, masking neoepitopes and the complete immunopeptidome^17^. As downregulation of antigen presentation is implicated in immune checkpoint blockade escape ^14^, methods to upregulate expression of the components of antigen presentation using chemotherapeutics or other agents are currently under investigation^18–24^. A number of groups have demonstrated that HLA expression and antigen presentation are transcriptionally silenced by repressive epigenetic marks^25–27^. Accordingly, epigenetic modifiers and immunotherapy are being explored as rational combination therapeutics for their efficacy at relatively non-toxic doses and ability to selectively reprogram cancer cells^23,28,29^. Moreover, DNA demethylating agents are actively being explored in preclinical models and in the clinic alongside checkpoint blockade inhibitors and other immunotherapies ^30–32^.

The oncogenic functions of EZH2 are being uncovered^33,34^. Activating mutations in the catalytic pocket of EZH2, such as those at Tyrosine 641 (Y641) cause excessive deposition of H3K27me3, which is associated with repressed transcription^27^. These mutations are common in DLBCL and are linked to tumor progression ^33,35^. Recent evidence also implicates EZH2 function in silencing anti-tumor immune responses ^23,24,29^. In line with these findings, EZH2 can directly recruit DNA methyltransferases to PRC2 target genes to further stabilize gene silencing ^36^. In DLBCL, over half of the *de novo* DNA methylation events overlap with PRC2 target genes, some of which are involved in the interferon-γ pathway ^37^. While manipulation of single epigenetic marks can reprogram transcription, epigenetic programs are redundant and highly coordinated ^38^ and therefore, targeting multiple epigenetic silencing pathways could more effectively activate expression of anti-tumor genes than single agent treatment. Therefore, understanding how combinatorial epigenetic treatment impacts potential responses to immunotherapy could have immediate high clinical impact.

Here, we explored the therapeutic potential of combining EZH2 inhibition using tazemetostat with DNA demethylation through decitabine in the presence of interferon gamma (IFNg). Both EZH2 inhibitors and DNA demethylating agents positively regulated antigen presentation in EZH2-mutated DLBCL cell lines and demonstrated combinatorial effects on transcriptional activation of antigen presentation from both MHC class I and II. The induced large increase in MHC surface expression of cryptic epitopes, especially in combination with IFNg enabled the comprehensive MS analysis of the normally heavily suppressed immunopeptidome of these DLBLCs. The drugs induced 10-fold to over 200-fold increases in total numbers of identified peptides presented by MHC class I, tracking with strong upregulation of MHC expression. The data demonstrated the feasibility of cryptic immunotherapy target identification in DLBCLs through MS. Among the many newly presented HLA ligands, several highly cancer-specific HLA ligands were identified that can serve as potential targets for immunotherapy design and combination therapies in DLBCL.

## Material and Methods

### Cell Culture

Cells were maintained in RPMI with penicillin and streptomycin supplemented with 10% heat-inactivated fetal bovine serum (FBS) and 5mM L-glutamine. All cells were maintained at 37C, 5% CO2. SUDHL-4 (A*02:01, B*15:01, C*03:04), DB (A*02:01, B*18:01, C*05:01), WSU-DLCL2, and Karpas 422 cells were from the Christian Steidl lab (British Columbia Cancer Research Centre) SUDHL-6 (A*02:01, A*23:01, B*15:01, B*49:01, C*03:03, C*07:01) and SUDHL-10 were provided by Dr. Anas Younes (Memorial Sloan Kettering Cancer Center). HLA typing was performed by the American Red Cross.

### Drug Treatments

Decitabine (Sellekchem, Cat. No S1200), tazemetostat (Sellekchem, Cat. No S7128) and interferon-γ (RnDsystems, 285-IF-100/CF) were administered *in vitro* using the same treatment schedule: Cells were treated with noted concentrations (Decitabine between 125 and 2000 nmol/ml, tazemetostat between 312.5 and 5000 nmol/ml, IFNg 1-100ng/ml) of each drug for 48hrs. Media was refreshed and new drug was added for an additional 48 hours.

### Flow Cytometry

Cells were treated as indicated in the respective sections. Cells were harvested, washed with PBS, and labeled in staining buffer (2% FBS, 0.1% sodium azide, in PBS) for 30 minutes with 1:400 dilution of FITC Mouse anti-Human HLA-A2 (clone BB7, Biolegend), APC anti-human HLA-A,B,C (clone W6/32, Biolegend), APC anti WT1-HLA-A*02:RMFPNAPYL (clone ESK1) antibodies. Clones Pr20 and ESK1 were generated and provided by Eureka Therapeutics Inc. and labeled with APC using the APC-Lightning Link®-Labeling Kit (Novusbio) according to manufacturer’s instructions. Cells were washed after incubation with staining buffer and analyzed using a Fortessa Flow Cytometer from BD Biosciences or a Guava flow cytometer from Millipore.

### Western Blot

Cells were treated as indicated in the respective sections. Total cell lysate was extracted using RIPA buffer and quantified using the DC protein assay (Bio-Rad). 15–30 μg of protein was loaded and run on 4%–12% SDS PAGE gels. After 1 hour block with 5% milk at room temperature, immunoblotting was performed using the following antibodies: anti-20s β5i (Enzo Life Science, BML-PW8845-0025), anti-20s β2i (Enzo Life Science, BML-PW8350-0025), and anti-20s β1i (Enzo Life Science, BML-PW8840-0025). Antibodies were probed at the manufacturer’s recommended dilution overnight at 4°C before a secondary antibody conjugated to HRP was used for imaging. Replicate samples were probed using the indicated antibodies when noted, or blots were stripped with Restore Western Blot Stripping Buffer (Thermo Fisher Scientific, 21063), re-blocked with 5% milk, and reprobed with an anti–GAPDH-HRP direct conjugated antibody (Cell Signaling Technology, 3683) as a loading control.

### Quantitative reverse-transcriptase PCR

Drug-treated cells were harvested and washed once with PBS. Cells were lysed in RLT buffer with Beta-mercaptoethanol and RNA was extracted using Qiagen RNEasy kit (Qiagen; #74134). Extracted RNA was then converted to cDNA using the one-step qSCRIPT cDNA solution. 5ng of isolated cDNA per sample were mixed with 1x target primer and 1x endogenous control primer in Perfecta master mix (Quantabio; #95118). Reactions were performed in the thermocycler. Primers used from Thermo Fisher are as follows : Hs00388675_m1; Human TAP1, Hs00241060 _m1; TAP2, Hs00984230_m1; Human B2M, Hs01058806_g1 Human HLA-A, Hs00818803_g1 Human HLA-B, Hs00740298_g1 Human HLA-C.

### Immunopurification of HLA ligands

HLA class I ligands (HLA-A, B and -C) and HLA class II ligands (HLA-DR) were isolated as described previously ^39^. In brief, 40 mg of Cyanogen bromide-activated-Sepharose 4B (Sigma-Aldrich, cat. # C9142) were activated with 1 mmol/L hydrochloric acid (Sigma-Aldrich, cat. # 320331) for 30 minutes. Subsequently, 0.5 mg of W6/32 antibody (Bio X Cell, BE0079; RRID: AB_1107730) or L243 antibody (Bio X Cell, BE0306; RRID: AB_2736986) was coupled to sepharose in the presence of binding buffer (150 mmol/L sodium chloride, 50 mmol/L sodium bicarbonate, pH 8.3; sodium chloride: Sigma-Aldrich, cat. # S9888, sodium bicarbonate: SigmaAldrich, cat. #S6014) for at least 2 hours at room temperature. Sepharose was blocked for 1 hour with glycine (Sigma-Aldrich, cat. # 410225). Columns were washed with PBS twice and equilibrated for 10 minutes. DB, SUDHL4 and SUDHL6 cells were treated with the indicated drugs. Cells (5×10^6^ to 1.5×10^7^) were harvested and washed three times in ice-cold sterile PBS (Media preparation facility MSKCC). Afterward, cells were lysed in 1 mL 1% CHAPS (Sigma-Aldrich, cat. # C3023) in PBS, supplemented with 1 tablet of protease inhibitors (Complete, cat. # 11836145001) for 1 hour at 4**°**C. This lysate was spun down for 1 hour at 20,000 g at 4**°**C. Supernatant was run over the affinity column through peristaltic pumps at 1 mL/minute overnight at 4**°**C. Affinity columns were washed with PBS for 15 minutes, run dry, and HLA complexes subsequently eluted five times with 200 mL 1% trifluoracetic acid (TFA, Sigma/Aldrich, cat. # 02031). For the separation of HLA ligands from their HLA complexes, tC18 columns (Sep-Pak tC18 1 cc VacCartridge, 100 mg Sorbent per Cartridge, 37–55 mm Particle Size, Waters, cat. # WAT036820) were prewashed with 80% acetonitrile (ACN, Sigma-Aldrich, cat. # 34998) in 0.1% TFA and equilibrated with two washes of 0.1% TFA. Samples were loaded, washed again with 0.1% TFA, and eluted in 400 mL 30% ACN in 0.1%TFA followed by 400 mL 40% ACN in 0.1%TFA, then 400 mL 50% ACN in 0.1%TFA. Sample volume was reduced by vacuum centrifugation for mass spectrometry analysis.

### LC/MS-MS analysis of HLA ligands

Samples were analyzed by a high-resolution/high-accuracy LC-MS/MS (Lumos Fusion, Thermo Fisher). Peptides were desalted using ZipTips (Sigma Millipore; cat. #ZTC18S008) according to the manufacturer’s instructions and concentrated using vacuum centrifugation prior to being separated using direct loading onto a packedin-emitter C18 column (75 mm ID/12 cm, 3 mm particles, Nikkyo Technos Co., Ltd). The gradient was delivered at 300 nL/ minute increasing linear from 2% Buffer B (0.1% formic acid in 80% acetonitrile)/98% Buffer A (0.1% formic acid) to 30% Buffer B/70% Buffer A, over 70 minutes. MS and MS/MS were operated at resolutions of 60,000 and 30,000, respectively. Only charge states 1, 2, and 3 were allowed. 1.6 Th was chosen as the isolation window and the collision energy was set at 30%. For MS/MS, the maximum injection time was 100 ms with an AGC of 50,000.

### Mass spectrometry data processing

Mass spectrometry (MS) data were processed using Byonic software ^40^ (version 2.7.84, Protein Metrics) through a custom-built computer server equipped with 4 Intel Xeon E5-4620 8-core CPUs operating at 2.2 GHz, and 512 GB physical memory (Exxact Corporation). Mass accuracy for MS1 was set to 6 ppm and to 20 ppm for MS2, respectively. Digestion specificity was defined as unspecific and only precursors with charges 1, 2, and 3, and up to 2 kDa were allowed. Protein FDR was disabled to allow complete assessment of potential peptide identifications. Oxidation of methionine, N-terminal acetylation, phosphorylation of serine, threonine, and tyrosine were set as variable modifications for all samples. All samples were searched against the UniProt Human Reviewed Database (20,349 entries, http://www.uniprot.org, downloaded June 2017). Peptides were selected with a minimal log prob value of 2 corresponding to p-values<0.01 for PSM in the given database and were HLA assigned by netMHCpan 4.0 ^41^ with a 2% rank cutoff.

### Statistical Methods

Experiments in figures 1, 2, 3, 4 and supplemental figures 1, 2, and 5 were analyzed using ANOVA followed by a post-hoc Tukey’s test to determine significance of individual groups. IFNg dose response experiments in figure 5 and supplemental figure 3 were analyzed using a student’s t test between control and treated conditions. Graphs and plots were made using R programming. Results from figure 5 and supplemental figure 5 and 6 were plotted using Graphpad Prism.

**Figure 1.**
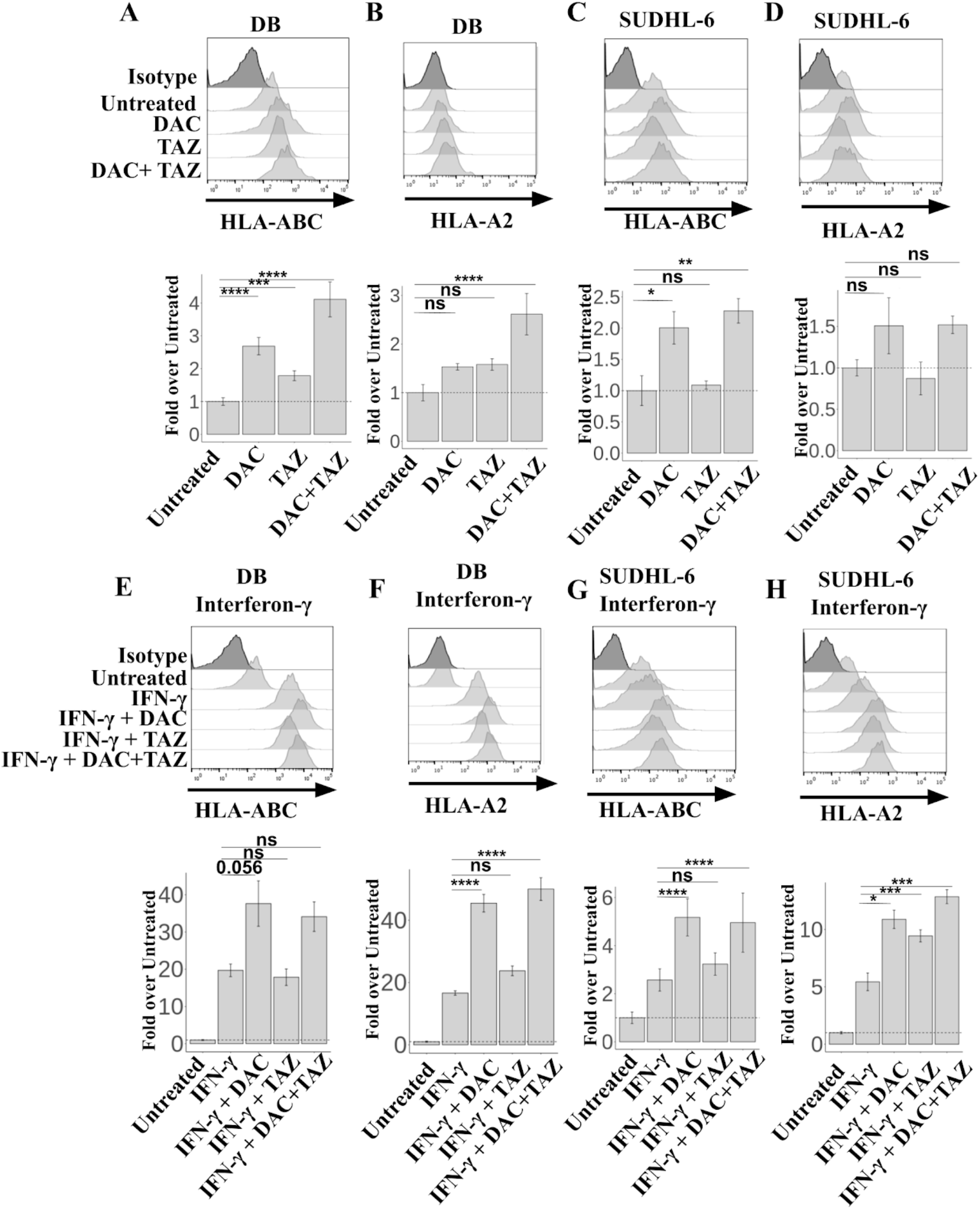
Decitabine and Tazemetostat upregulated HLA protein in DLBCL cell lines. (A-D) Cells were treated with indicated 125nM Decitabine, or 1uM Tazemetostat. (A-B) DB and (C-D) SUDHL-6 cells were assayed for HLA-ABC (A+C) and HLA-A-02 (B+D) expression by flow cytometry. (E-H) Cells were treated as in (A-D) alongside 100ng/mL IFNg. ANOVA using either untreated (A-D) or IFNg alone (E-H) as control was performed, followed by a post-hoc Tukey’s test for individual experimental groups. N=2 technical replicates per 2 biological replicates. * p<.05, ** p<.01, *** p<.001, **** p<.0001.

**Figure 2.**
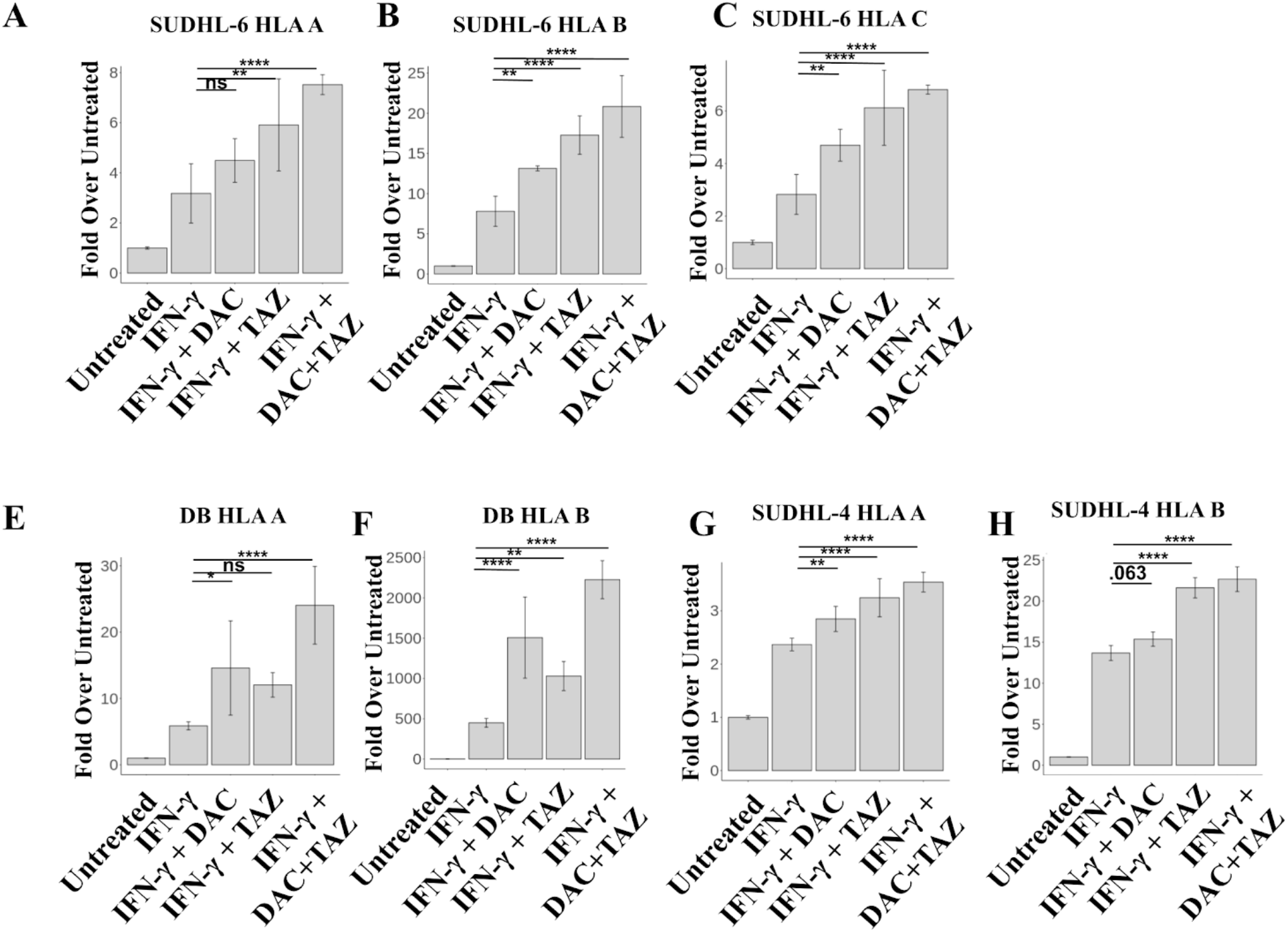
Decitabine and Tazemetostat activated transcription of HLA alleles on DLBCL cell lines. (A-D) SUDHL-6 and (E-H) DB cells were treated with indicated drugs. Decitabine was given at 50nM, Tazemetostat at 1uM, and IFNg at 10ng/mL. Graph of fold-change in transcript to untreated for each indicated gene (A+E) *HLA* (B+F) *B2M* (C+G) *Tap1* (D+H) *Tap2*. ANOVA using either untreated (A-D) or IFNg alone (E-H) as control was performed, followed by a post-hoc tukey’s test for individual experimental groups. N=3 technical replicates per 2 biological replicates. * p<.05, ** p<.01, *** p<.001, **** p<.0001.

**Figure 3.**
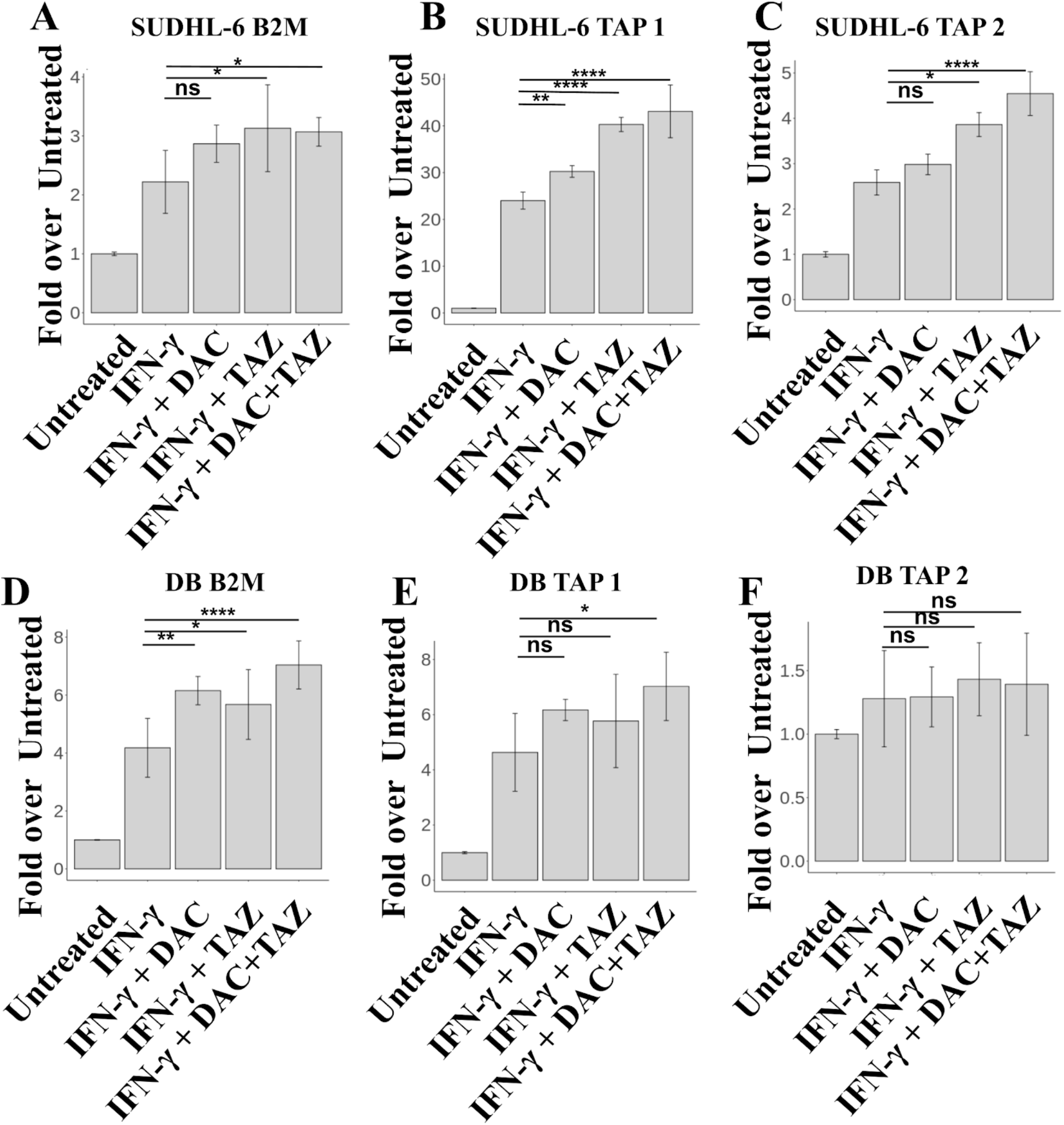
Decitabine and Tazemetostat activated transcription of antigen presentation genes in DLBCL cell lines. (A-D) SUDHL-6 and (E-H) DB cells were treated with indicated drugs. Decitabine was given at 50nM, Tazemetostat at 1uM, and IFNg at 10ng/mL. Graph of fold-change in transcript to untreated for each indicated gene (A+E) *HLA* (B+F) *B2M* (C+G) *Tap1* (D+H) *Tap2*. N=3 technical replicates per 2 biological replicates. * p<.05, ** p<.01, *** p<.001, **** p<.0001.

**Figure 4.**
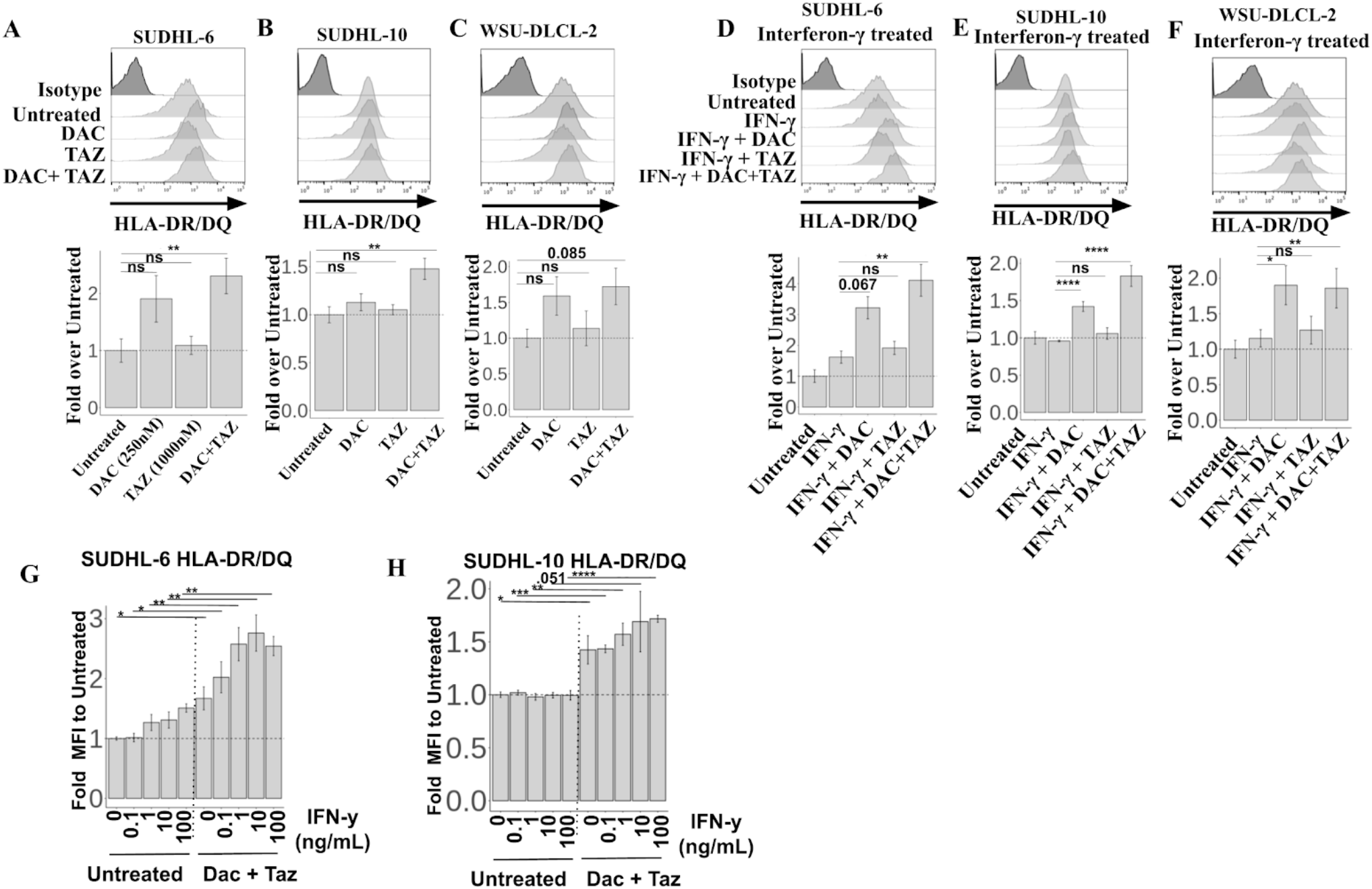
Decitabine upregulated HLA Class II in DLBCL cell lines. (A-C) Cells were treated with indicated 125nM decitabine, or 1uM Tazemetostat for (A) SUDHL-6 (B) SUDHL-10 (C) WSU-DLCL-2 cell lines, and assayed for expression of HLA-DR/DQ. (D-F) Cells were treated as in (A-C) alongside 100ng/mL IFNg. (G-H) Serial dilutions of IFNg were performed in the presence or absence of decitabine and tazemetostat for (G) SUDHL-6 and (H) SUDHL-10 cell lines. N=2 technical replicates per 2 biological replicates. * p<.05, ** p<.01, *** p<.001, **** p<.0001.

**Figure 5.**
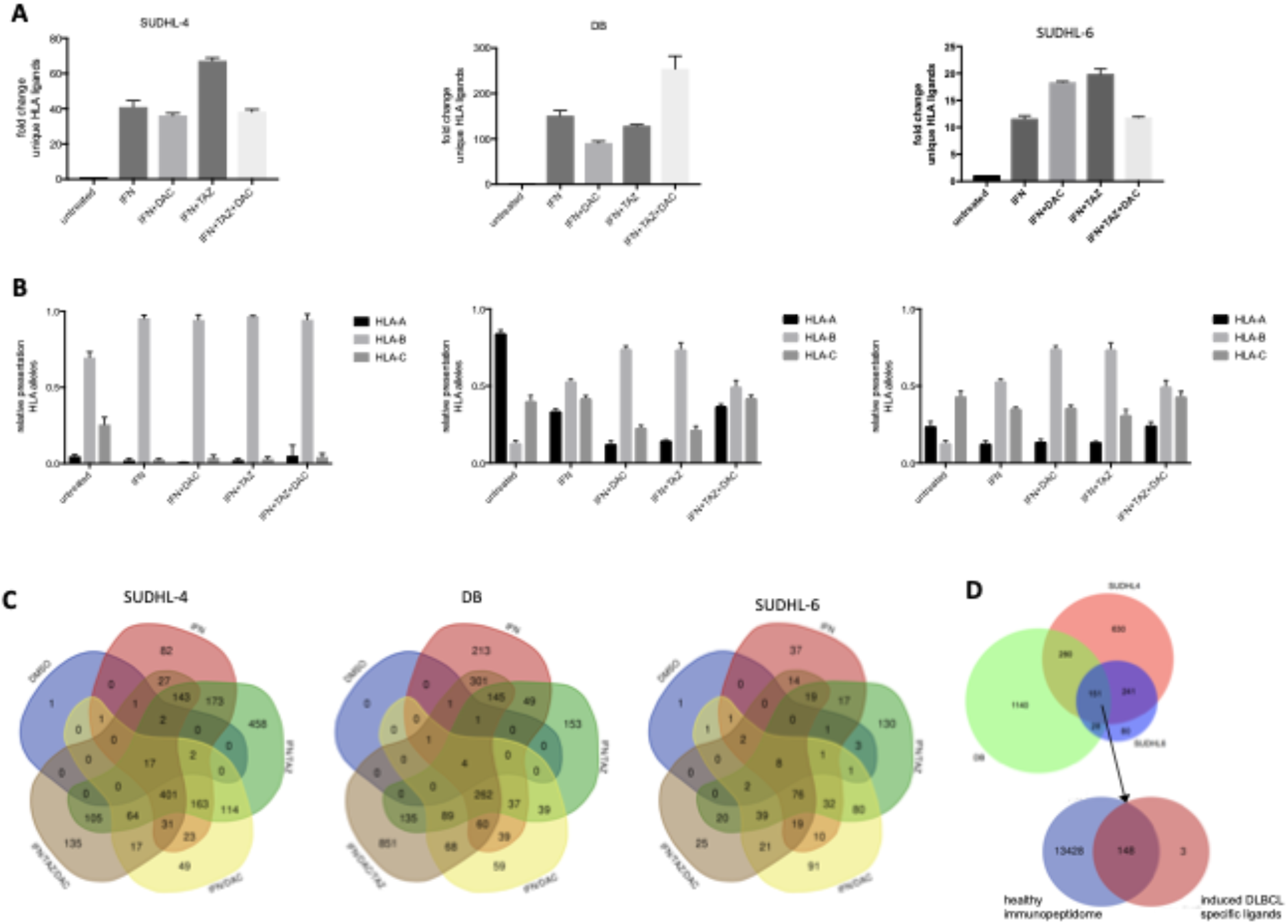
Epigenetic drug treatment in presence of IFNg unmasked the immunopeptidome of DLBCL cell lines. (A) Cells were treated with 125nM decitabine, or 1uM tazemetostat or the combination of both in presence of 100 ng/mL IFNg. Fold change of unique identifications of HLA ligands relative to untreated cells is depicted. Error bars indicate mean plus SD. Experiments were performed in duplicates. (B) Relative distribution of HLA alleles after assignment to their respective alleles through netMHCpan 4.0. (C) Overlap analysis of all peptides by cell line and respective treatment condition. (D) Overlap of source proteins for HLA ligands shared between the cell lines SUDHL-4, DB, SUDHL-6 (top), Overlap of 151 source proteins from overlap at the top were matched with 13,428 source proteins of the HLA class I ligandome from healthy donors as published by Marcu et al.^43^

## Results

### Decitabine and Tazemetostat improved IFNg responsive HLA upregulation in EZH2 mutated DLBCL cell lines

To understand how decitabine and tazemetostat might impact antigen presentation, we first measured HLA I cell surface expression in EZH2-mutated DLBCL cell lines via flow cytometry. We selected cell lines to be HLA-A*02:01 positive as it represents the most common HLA haplotype in the western hemisphere and therefore is of broad interest. Both DB and SUDHL-6 treated DLBCL cell lines variably upregulated HLA I and HLA-A*02 cell surface expression when treated with 1uM tazemetostat (TAZ), a highly specific EZH2 inhibitor, 125 nM decitabine (DAC), a potent DNA demethylating agent, or a combination of both (Figure 1A-D). Interestingly, decitabine led to a more robust upregulation of HLA as monotherapy compared to tazemetostat. Trends toward increased HLA I expression were also observed for the EZH2 mutant DLBCL cell lines WSU-DLCL2, SUDHL-4, SUDHL-10, but not Karpas 422 (Supplemental Figure 1). When DLBCL cell lines were treated with interferon-γ in addition to the aforementioned treatment schema, there was a cooperative increase in total HLA I expression reaching over 50-fold for DB cells, as well as 5-fold upregulation for SUDHL-6 and over 50-fold for DB cells when HLA-A*02 alone was assessed (Figure 1E-H). Similarly, modest trends in increased HLA expression were seen in WSU-DLCL2, SUDHL-4, SUDHL-10, but not Karpas 422 (Supplemental Figure 2).

To determine whether TAZ synergizes with DAC in HLA I upregulation, we performed a dose titration of DAC in the presence of 625nM TAZ and 100ng/mL IFNg and tested for HLA-A*02, as most profound changes were seen for this allele in the previous experiment. Indeed, a constant concentration of TAZ shifted the expression of HLA-A*02 at each DAC concentration tested (Supplemental Figure 3A). Similarly, a shift in expression was seen by a TAZ titration if the concentration of DAC was kept constant at 250nM (Supplemental Figure 3B). Additionally, TAZ alone showed a dose dependent increase in HLA expression in SUDHL-4 and DB cells, in the presence or absence of IFNg (Supplemental Figure 4). Finally, to demonstrate that DAC and TAZ are able to increase IFNg responsiveness, we performed 10-fold dilutions of IFNg, in the presence of 125nM DAC and 1uM Taz. DB and SUDHL-6 cells showed significantly higher responses to IFNg reaching again 50-fold upregulation for DB cells and increases up to 6-fold for SUDHL-6 cells when treated with epigenetic modifiers (Supplemental Figure 3C-D). The enhanced HLA expression in triple treated cells demonstrates potential compensatory mechanisms utilized by the cancer cells to suppress antigen presentation and interferon-γ responsiveness, that can be rewired through epigenetic treatment.

### Decitabine and Tazemetostat modulated individual HLA alleles independently and upregulated the transcription of antigen presentation machinery in DLBCL

Given the general upregulation of HLA after treatment with TAZ or DAC, we further evaluated the impact of these drugs on transcription of each HLA allele individually as well as their impact on other components of the antigen presentation machinery using quantitative reverse-transcriptase PCR (qRT-PCR). SUDHL-6 cells showed a progressive increase in transcription of each allele, which was most prominent in the triple treatment group. Additionally, confirming prior reports, IFNg upregulated HLA-B transcription more profoundly than HLA-A and -C (Figure 2A-C)^42^. DB cells similarly showed a progressive increase in expression of HLA-A and -B loci when treated in combination with IFNg, DAC and TAZ (Figure 2E-F). DAC had a stronger impact on HLA upregulation compared to TAZ. Remarkably, HLA-B expression was increased 500-fold with IFNg treatment, which could be improved to about 1500-fold with combination DAC, 1000-fold with combination TAZ, and 2200-fold in the triple treatment group across two independent biological replicates. For the DB cell line, HLA-C levels were undetectable. Finally, SUDHL-4 cells also showed significant upregulation of HLA-A and -B in combination epigenetic treatment (Figure 2G-H). As with DB, SUDHL-4 cells showed no expression of HLA-C transcript in multiple experimental repeats.

Given large upregulations of the different HLA I alleles during drug treatment, we assessed upregulation of antigen presentation machinery β2-microglobulin (β2M), and Transporter associated with Antigen Processing (TAP) 1 and 2. Although less striking than HLA expression, β2M, TAP1 and TAP2 were significantly increased in SUDHL-6, and β2M and TAP1 were upregulated in DB cells across two independent biological replicates, when IFNg-treated cells were also treated with DAC or TAZ (Figure 3). Similar, but less profound, upregulation of antigen presentation was also seen in SUDHL-4 cells (Supplemental Figure 5). Yet, overall EZH2 and DNA methylation inhibition upregulated transcripts involved in antigen processing and presenting at very high levels.

### Decitabine Upregulates HLA Class II Expression in DLBCL

To complete the analysis of proteins primarily involved in antigen presentation we turned to HLA class II as it is also expressed highly on DLBCLs in line with their B cell origin. The overall limited expression of HLA class II throughout the body also makes class II presented peptides suitable immunotherapy targets. Therefore, we assessed the impact of DAC and TAZ on expression of HLA Class II. TAZ had no significant effect on HLA DR/DQ expression. However, DAC treatment trended towards upregulation of HLA DR/DQ, while the combination led to significant upregulation in SUDHL-6 and SUDHL-10, and a trend in WSU-DLCL2 cells (Figure 4A-C). Class II upregulation was further enhanced by addition of IFNg to the DAC and TAZ treated cells (Figure 4D-F). To determine if DAC and TAZ lowered the threshold for IFN-g mediated Class II upregulation, we titrated IFNg in the presence of 125nM DAC and 1uM TAZ. As seen with HLA Class I expression, HLA DR/DQ was significantly upregulated by DAC and TAZ across each IFNg concentration (Figure 4G, H).

### Epigenetic drug treatment in combination with IFNg allowed systematic analysis of the DLBCL immunopeptidome and identified disease-specific HLA ligands

Given the extensive increases in HLA class I and class II expression, antigen presentation machinery, and individual ligand presentation, which followed the treatment with epigenetic modifiers in combination with IFNg, we wanted to systematically investigate the changes and potential emergence of new peptides that are presented on the cell surface. Using immunoprecipitation with HLA-A,B,C and HLA-DR specific antibodies, separation of the bound peptides and subsequent mass spectrometry we identified increases in the numbers of unique HLA ligands similar to the fold changes seen in RT-qPCR and by flow cytometry for HLA levels. Strikingly, this method, which usually robustly identifies thousands of different HLA ligands in a single sample, detected few (between 7 and 19) unique peptides in the three untreated cell lines SUDHL4, DB and SUDHL6, which resembles the profound downregulation of HLA levels in these cell lines. In contrast, when cell lines were treated with IFNg and the previously used schema of DAC and TAZ, SUDHL6 cells presented over 400 unique HLA ligands, SUDHL-4 cells over 1,500 unique HLA ligands, and DB cells over 2,000 unique HLA ligands using the most effective treatment condition. This number of ligands is similar to numbers of peptides identified on healthy B cells if similar cell numbers were used (Supplemental Figure 6 A-C). Thus, this corresponds to an upregulation of HLA ligand presentation in the range of 10-20 fold for SUDHL-6, 40-70 fold for SUDHL-4, and 100-250 fold for DB cells (Figure 5A). Though the combination of TAZ and IFN induced the strongest changes in SUDHL-4 and SUDHL-6 cells, no clear drug combination induced the most unique peptides. In all three cell lines, the addition of TAZ or TAZ in combination with DAC showed additive effects over the IFNg treatment alone.

Next, we determined the allelic distribution of these newly presented HLA ligands. Therefore, we used netMHCpan 4.0 to assign each peptide to one of the expressed HLA alleles and then looked at the fraction of peptides presented on each allele. First, all cell lines demonstrated the known preferences for peptide presentation on HLA-B alleles after IFNg treatment. Interestingly, all epigenetic monotherapies (TAZ or DAC) enhanced this effect. In contrast, the combination of both modifiers led to distribution patterns identical to IFNg treatment alone (Figure 5B).

Furthermore, the HLA ligand overlaps between the five treatment conditions in each cell line demonstrated that for each single treatment condition, a relevant proportion of peptides that was unique to this treatment group was detectable (Figure 5C). This fraction could be as little as 8% (82/1022 peptides), e.g. in the IFNg treated SUDHL-4 cells, but these fractions could be as high as 29-30% as seen for the IFN/TAZ treated subgroups in SUDHL-4 (458/1555 peptides) and SUDHL-6 (130/431 peptides). The triple-treated condition in DB cells displayed HLA ligands of which 44% (851/1918 peptides) were unique to that treatment condition. No consistent pattern could be observed regarding which epigenetic treatment led to the strongest improvement, though it was evident that these drugs could lead to substantial changes in the immunopeptidome resulting in presentation of many HLA I ligands previously not displayed.

Finally, we investigated the origin of peptides that arose after different treatment conditions. First, to analyze if the presented peptides correlated with the phenotype of a DLBCL, we specifically looked for HLA ligands derived from proteins that serve as histological biomarkers in the diagnosis of DLBCL or that are associated with relevant pathobiology of the disease (BCL-2, BCL-6, MYC, MYD88, CD79A/B, CREBBP, PAX5 and EZH2). HLA ligands could be identified for four of these eight proteins. For bcl-6 and CD79A/B multiple HLA ligands on different HLA alleles were detectable after drug treatment, whereas for the CREBBP and PAX-5 proteins, only one HLA ligand per protein was found (Table 1, top).

**Table 1.**
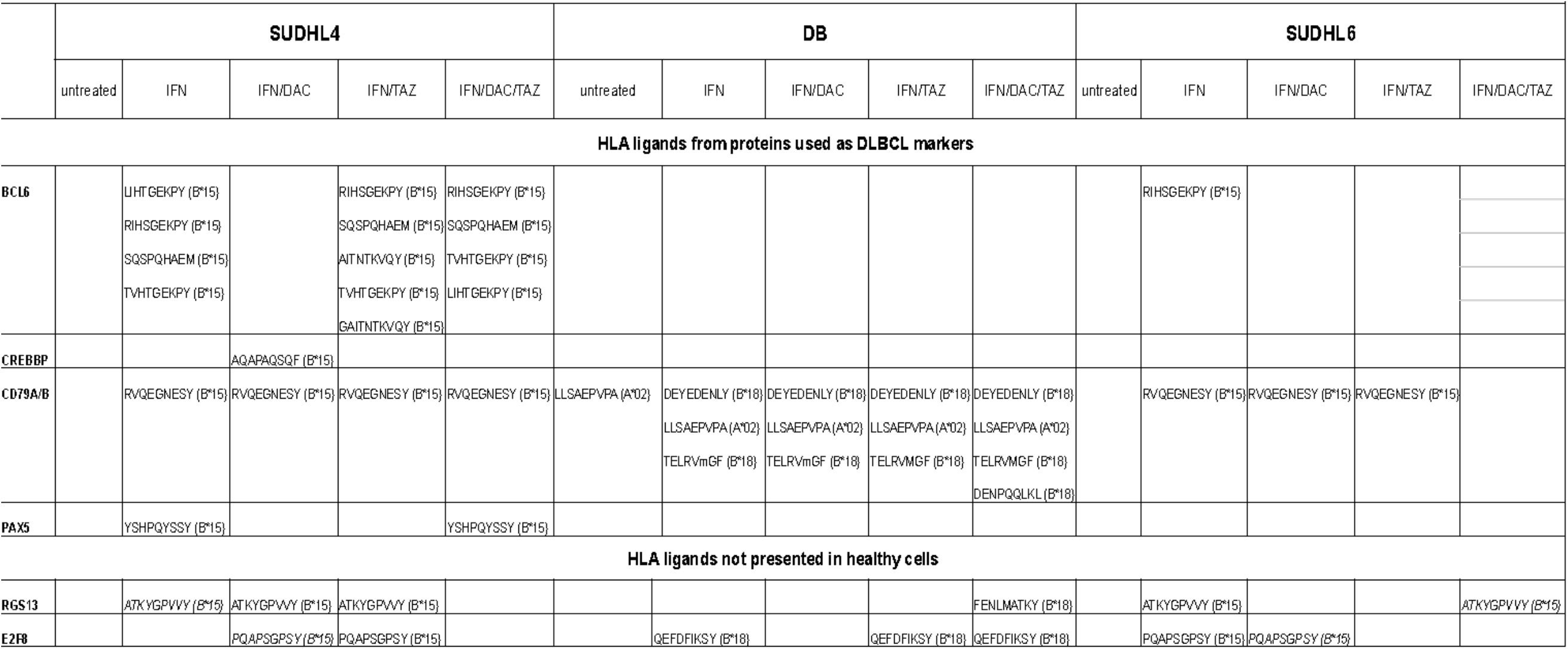
HLA ligands identified from DLBCL markers and disease specific HLAligands

More importantly, to identify potential immunotherapy targets, we looked for HLA ligands that met the following criteria: 1) peptides derived from proteins that were identified in all three cell lines and 2) peptides from proteins that have not been reported in the immunopeptidome of healthy donors. Therefore, we overlapped the source proteins from which the HLA ligands were derived, from all three cell lines and identified 151 proteins shared between all three lines. At this point we focused only on the protein, but not the HLA level to account for the different HLA alleles present in these cell lines. We then took these 151 proteins and matched them against 13,576 proteins, which were recently reported by Marcu et al. to be the source of HLA ligands in healthy tissues^43^. From this analysis only three proteins were identified to be unique to the DLBCL group (Figure 5D). One protein, an HLA allele subtype, could also be annotated to proteins included in the 13,576 proteins after further analysis and therefore was determined to be a false positive. The remaining two proteins, *regulator of G protein signaling 13* (RGS13) and *E2F transcription factor 8* (E2F8), were highly tumor-specific, as no HLA class I ligands from these proteins have been reported on healthy tissues. RGS13 derived peptides were identified in other malignant samples and E2F8 derived HLA ligands were exclusively found in transformed B cells, according to IEDB database ^44^. Moreover, the identified HLA ligands from RGS13 and E2F8 in these cell lines were detectable in many different treatment replicates; in case of ATKYGPVVY (B*15) and PQAPSGPSY (B*15) the epitope was shared between the cell lines SUDHL-4 and SUDHL-6, which renders these HLA ligands promising targets for immunotherapy.

## Discussion

The immunopeptidome of many hematological malignancies has been described in detail and provides a valuable source for possible targets of T cell immunotherapy ^11–13,45^. In contrast, little immunopeptidome data are available for DLBCL, most likely due to a strong HLA downregulation, which has been reported especially for the EZH2-mutated subgroup ^23^. Identifying possible targets in DLBCL is important as immunotherapies are often powerful therapies with relatively low toxicity.

Here, we demonstrated how EZH2 inhibitors and DNA demethylation in presence of IFNg can overcome the immune evasion mechanisms of DLBCL, leading to robust unmasking of novel, potentially targetable HLA ligands. First, these drug combinations led to substantial increases of HLA class I surface expression with increases up to 50-fold by flow cytometry, which were mediated to a large extent by IFNg, but which were clearly enhanced by the addition of TAZ or DAC or the combination of both drugs. The therapeutic combination of DNA demethylating agents with selective EZH2 inhibitors is supported by evidence of direct, physical links between EZH2 and DNA methyltransferases. EZH2 recruits DNA methyltransferases to PRC2 gene loci for further stabilization of the repressive epigenetic program ^36^. Therefore, genes that are initially silenced by overactive EZH2 function in DLBCL may be resistant to upregulation by EZH2 inhibitors if DNA methylation is redundantly silencing the locus. Indeed, over half of the *de novo* DNA methylation events between healthy B cells and B cell lymphoma are at PRC2 target sites^37^.

Moreover, when we investigated the allele specific HLA upregulation we discovered that HLA-C alleles were not detectable in DB and SUDHL-4 cells. Because these cell lines were already homozygous for all three HLA class I alleles this loss of HLA-C significantly reduces the number of HLA alleles that were able to present a diverse immunopeptidome to two alleles. Moreover, when we quantitatively examined the peptides presented by these cell lines, few HLA ligands were assigned to HLA-A*02 in SUDHL-4, rendering this cell line functionally nearly mono-allelic, which illustrates again the profound immunosuppressive mechanisms in DLBCL, and another strategy for immune evasion in lymphoma.

Finally, the drug-induced unmasking of HLA ligands in the cell lines we investigated allowed us, for the first time, to conduct a systematic analysis of HLA ligands present in DLBCL, which led to the detection of HLA I ligands from RGS13 and E2F8, which had not been observed in healthy tissues. Interestingly, ATKYGPVVY from RGS13 has not been described in any other cancer sample, demonstrating that truly novel HLA ligands can arise after these drug treatments. As two of the HLA ligands from HLA-B*15 (ATKYGPVVY and PQAPSGPSY) from RGS13 and E2F8, respectively were found on both cell lines expressing this allele (SUDHL4 and SUDHL6), we also demonstrated the feasibility of identifying tumor-specific *shared* antigens, which could be targeted independently of their immunogenicity by TCR-based agents or TCR mimic antibodies.

## Acknowledgements

We thank Alex Kentsis for access to the Byonic Software and Henrik Molina from the Proteome Resource Center at The Rockefeller University for the performance of all LC/MS-MS experiments. We thank Steven Josefowicz for guidance during this project. This study was supported by the Leukemia and Lymphoma Society; NIH P30 CA008748, R01 CA55349, P01 CA23766 and R35 CA241894; and Tudor Funds. MGK is supported by the German Research Foundation (DFG) with an individual Research Grant KL3118/1-1.

## Authorship contributions

C.M.B., S.S.M., Z.E.H.A., and M.G.K., performed and analyzed experiments. C.M.B., M.G.K., K.T., A.D, R.S.G., and D.A.S. designed experiments. C.M.B., and M.G.K. wrote the original draft of the manuscript. T.D., S.J., C.S., M.G.K and D.A.S. supervised the project. D.A.S. provided funding and edited the manuscript. All authors reviewed and contributed to the manuscript.

## Supplementary Information for

**Supplemental Figure 1.**
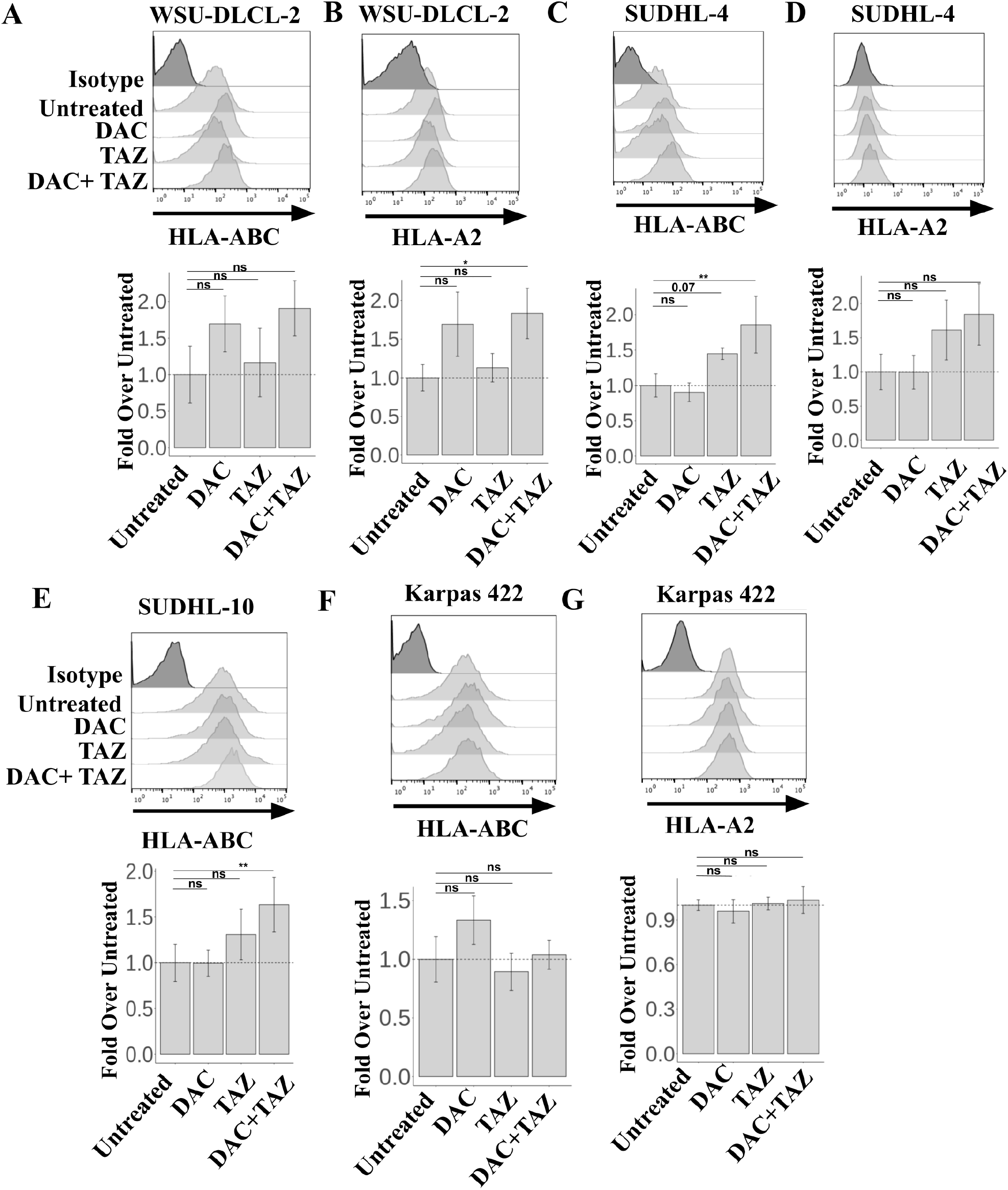
Decitabine and Tazemetostat upregulated HLA in WSU-DLCL2, SUDHL-4, SUDHL-10, but not Karpas 422 cell lines. (A-G) Cells were treated with indicated 125nM Decitabine, or 1uM Tazemetostat. (A-B) WSU-DLCL2, (C-D) SUDHL-4, (E) SUDHL-10, and (F-G) Karpas 422 cells were assayed for HLA-ABC (A,C,E,F) and HLA-A-02 (B,D,G) expression by flow cytometry. ANOVA using untreated as control was performed, followed by a post-hoc tukey’s test for individual experimental groups. N=2 technical replicates per 2 biological replicates. * p<.05, ** p<.01, *** p<.001, **** p<.0001.

**Supplemental Figure 2.**
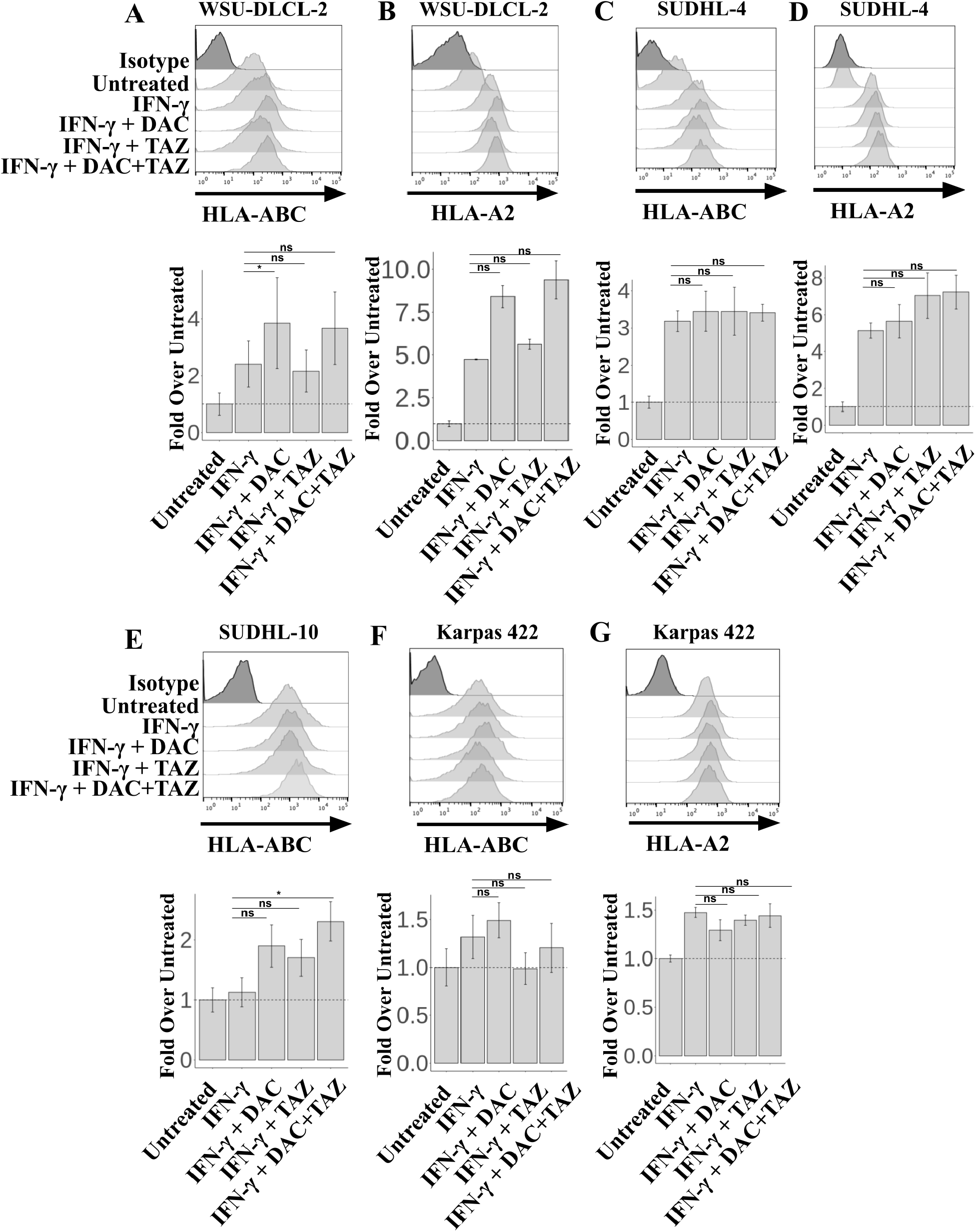
Decitabine and Tazemetostat on IFNg-mediated HLA upregulation in WSU-DLCL2, SUDHL-4, SUDHL-10, and Karpas 422 cell lines. (A-G) Cells were treated with indicated 125nM Decitabine, or 1uM Tazemetostat alongside 10ng/mL IFNg. (A-B) WSU-DLCL2, (C-D) SUDHL-4, (E) SUDHL-10, and (F-G) Karpas 422 cells were assayed for HLA-ABC (A,C,E,F) and HLA-A-02 (B,D,G) expression by flow cytometry. ANOVA using IFNg as control was performed, followed by a post-hoc tukey’s test for individual experimental groups. N=2 technical replicates per 2 biological replicates. * p<.05, ** p<.01, *** p<.001, **** p<.0001.

**Supplemental Figure 3.**
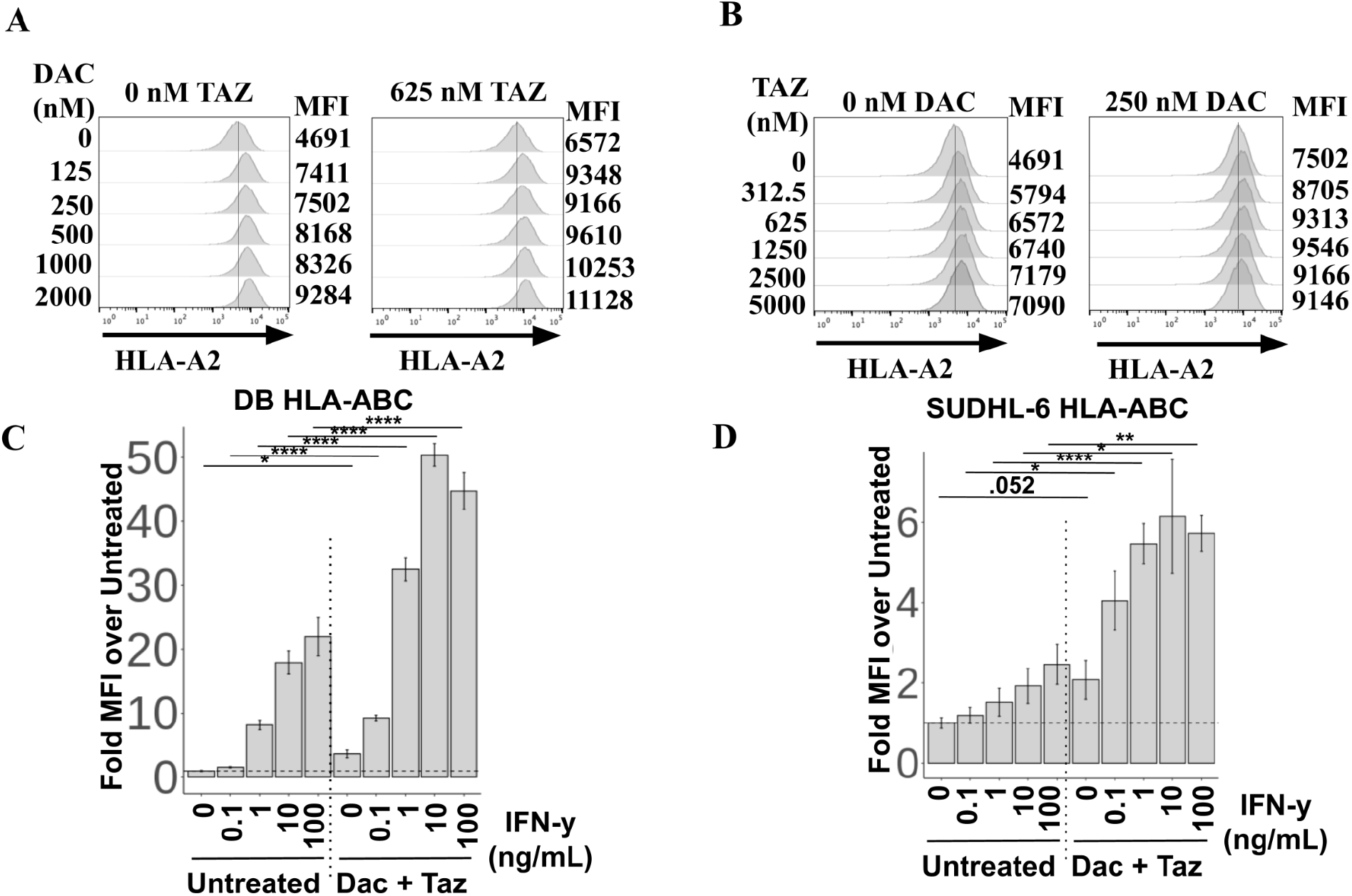
Decitabine and tazemetostat both regulated HLA expression in DLBCL in a dose responsive manner. (A-B) SUDHL-4 cells were treated with 100 ng/mL Interferon-γ. (A) Dose responses of decitabine were performed in the absence (left) or presence (right) of 625nM tazemetostat. (B) Dose responses of tazemetostat were performed in the absence (left) or presence (right) of 250nM decitabine. (C-D) A dose response of IFN-g was performed plus or minus decitabine and tazemetostat on (C) DB or (D) SUDHL-6 cells. A student’s T test was performed between untreated and Dac+ Taz at each IFNg concentration. * p<.05, ** p<.01, *** p<.001, **** p<.0001.

**Supplemental Figure 4.**
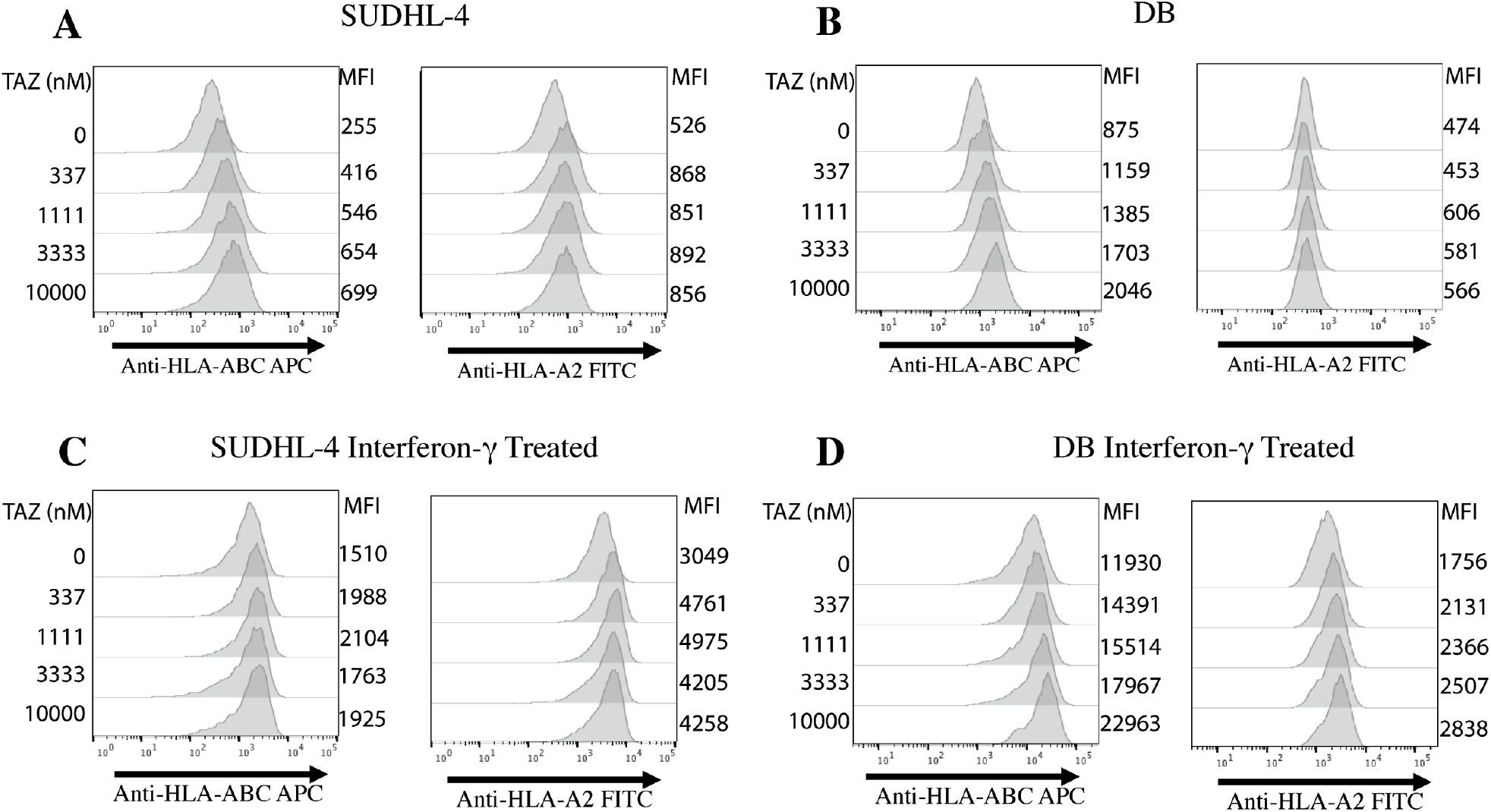
Tazemetostat upregulated HLA expression on DLBCL in a dose dependent manner. (A) SUDHL-4 cells and (B) DB cells were treated with indicated concentrations of tazemetostat. Left panel demonstrates the HLA-ABC expression and the right panel shows the HLA-A2 expression. (C-D) Cells were treated as in (A-B) in addition to 100ng/mL IFNg.

**Supplemental Figure 5.**
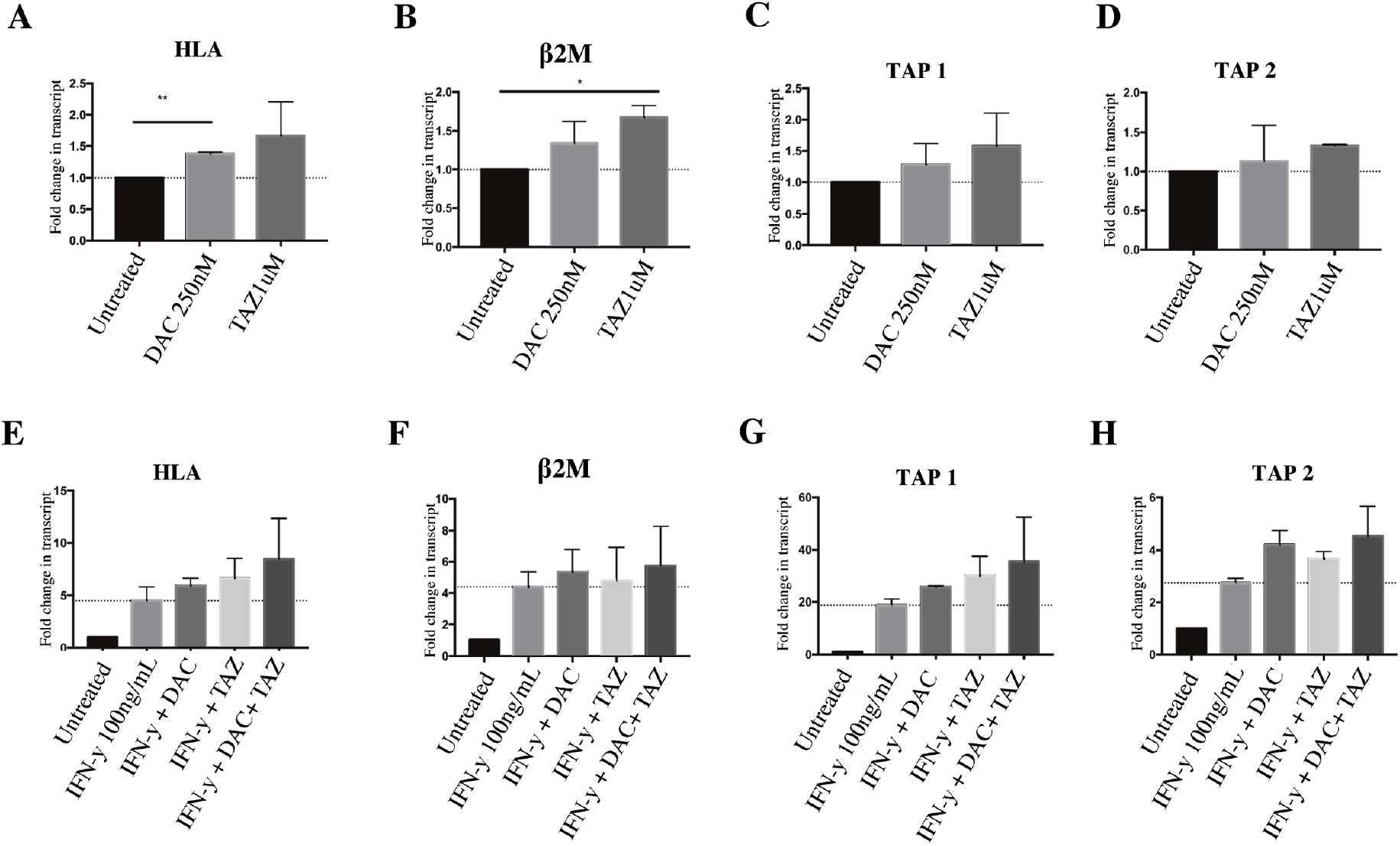
SUDHL-4 cells showed modest increases in antigen presentation transcript. (A-D) SUDHL-4 cells were treated with either 250nM Dac, 1uM Taz and transcript levels of (A) HLA-A, (B) beta-2-microglobulin, (C) TAP 1, and (D) TAP 2 were assessed. (E-F) Cells were treated as in (A-D) in addition to IFNg. ANOVA using (A-D) untreated or (E-F) IFNg as control was performed, followed by a post-hoc tukey’s test for individual experimental groups. N=2 technical replicates per 2 biological replicates. * p<.05, ** p<.01, *** p<.001, **** p<.0001.

**Supplemental Figure 6.**
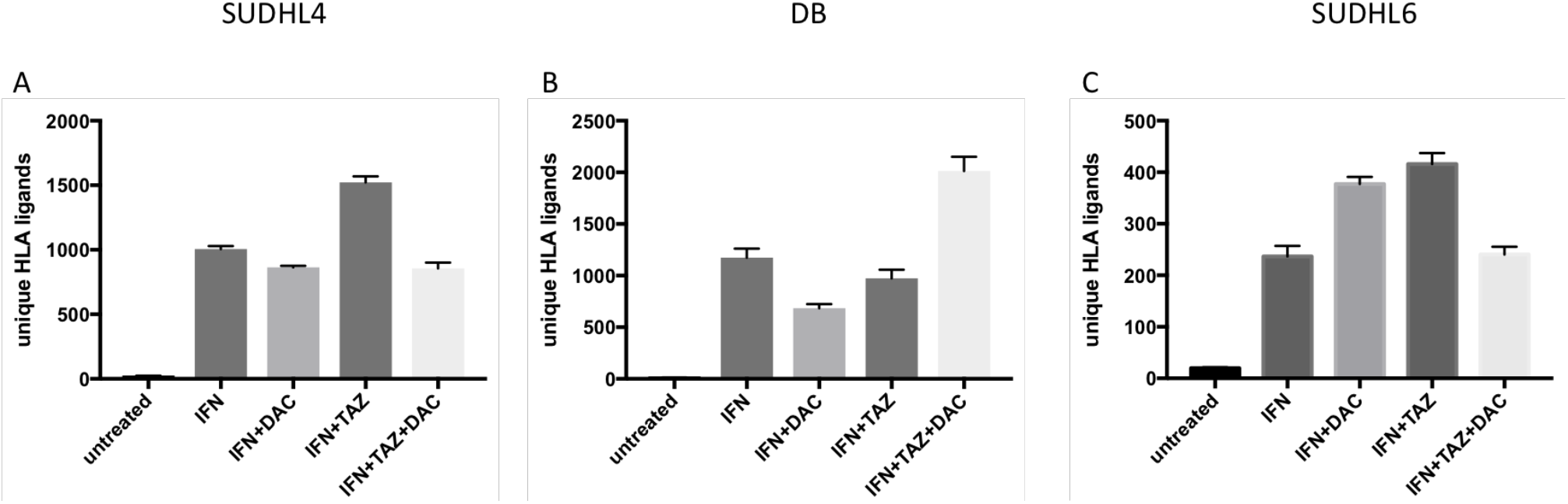
Epigenetic drug treatment in combination with IFNg induced presentation of HLA ligands in DLBCL cell lines. Cells were treated with 125 nMdecitabine, 1uMtazemetostatand 100 ug/ml IFNg. Treatments were applied to (A) SUDHL-4, (B) DB and (C) SUDHL-6 cells. Results show unique HLA ligands identified by mass spectrometry after assignment by netMHCpan4.0. Results are shown for 2 biological replicates. Error bars indicate mean with standard deviation.

